# Causal explanation of individual differences in human sensorimotor memory formation

**DOI:** 10.1101/255091

**Authors:** Pierre Petitet, Jill X. O’Reilly, Ana M. Gonçalves, Piergiorgio Salvan, Shigeru Kitazawa, Heidi Johansen-Berg, Jacinta O’Shea

**Affiliations:** Wellcome Centre for Integrative Neuroimaging (WIN), Oxford Centre for Functional MRI of the Brain (FMRIB), Nuffield Department of Clinical Neurosciences (NDCN), University of Oxford, John Radcliffe Hospital, Headington, Oxford, U.K.; Donders Institute for Brain, Cognition and Behaviour, Radboud University, Nijmegen, Netherlands; Dynamic Brain Network Laboratory, Graduate School of Frontier Biosciences, Osaka University, Osaka, Japan; Department of Brain Physiology, Graduate School of Medicine, Osaka University, Osaka, Japan; Center for Information and Neural Networks, Osaka University, Osaka, Japan

## Abstract

Sensorimotor cortex mediates the formation of adaptation memory. Individuals differ in the rate at which they acquire, retain, and generalize adaptation. We present a mechanistic explanation of the neurochemical and computational causes of this variation in humans. Neuroimaging identified structural, functional and neurochemical covariates of a computational parameter that determines memory persistence. To establish causality, we increased sensorimotor cortex excitability during adaptation, using transcranial direct current stimulation. As predicted, this increased retention. Inter-individual variance in the stimulation-induced E:I increase predicted the computational change, which predicted the memory gain. These relations did not hold, and memory was unchanged, with stimulation applied before adaptation. This cognitive state dependent effect was modulated by the BDNF val66met genetic polymorphism. Memory was enhanced by stimulation in Val/Val carriers only, implicating a mechanistic role for activity-dependent BDNF secretion. Sensorimotor cortex E:I causally determines the time constant of memory persistence, explaining phenotypic variation in adaptation decay.

## Introduction

Individuals differ – an explanatory challenge. Here we exploit this variation to test a causal mechanistic hypothesis about how sensorimotor memories in human brain are formed, retained and enhanced. Primary motor cortex (M1) has been identified as a critical region for the maintenance of newly acquired visuomotor maps following sensorimotor adaptation in both primates (Wise et al. 1998; Li et al. 2001; Paz et al. 2003; Paz et al. 2005; Inoue et al. 2016) and humans (Hadipour-Niktarash et al. 2007; Hunter et al. 2009; Galea et al. 2010; Landi et al. 2011; Leow et al. 2016; O’Shea et al. 2017). We previously demonstrated that left M1 anodal transcranial direct current stimulation (a-tDCS) applied during adaptation to a 10-degree rightward displacement of the visual field (prism adaptation, PA) enhances consolidation of the prism after-effect (AE) in the healthy brain (O’Shea et al. 2017). Notably, this stimulation-induced memory enhancement was 1) anatomically specific (it was not observed with left parietal or right cerebellar tDCS), 2) polarity specific (it was not observed with left M1 cathodal tDCS), and 3) cognitive state dependent (it was not observed when a-tDCS was applied before as opposed to during PA). Here, we interrogate the computational and neural bases of this effect.

Since adaptation memory was enhanced only with M1 excitatory stimulation during PA, the interaction with information processing in M1 was critical. What computation did stimulation alter to cause memory change? Computational theories of motor control posit that when a perturbation (such as an optical shift) causes reaching errors, the brain forms an estimate of the perturbation, used to counteract the disturbance and correct movement accuracy (Shadmehr et al. 2010; Franklin and Wolpert 2011; Wolpert et al. 2011; Petitet et al. 2017). For future movements to also be accurate, this estimate should not only include a representation of the magnitude of the perturbation (e.g. by how much do I need to correct?) but also of its temporal dynamics (e.g. for how long is the perturbation likely to prevail?). The latter is known as a temporal credit assignment problem. For retention to be optimal, it should match the internal estimate of the perturbation timescale, i.e. the solution to the temporal credit assignment (Körding et al. 2007). For example, if the disturbance is believed to be stable, adaptation should be retained; if transient, it should decay. Thus we reasoned that it might be this computation, whose implementation is likely to involve M1, that interacts with a-tDCS such that memory is enhanced.

If that is true, what features of M1 physiology are likely to be involved in this computation? Primary motor cortex a-tDCS is known to locally increase the excitation:inhibition (E:I) ratio via a reduction of gamma-aminobutyric acid (GABA) concentration (Stagg et al. 2009; Kim et al. 2014; Bachtiar et al. 2015; Antonenko et al. 2017). In our previous study, the magnitude of the stimulation-induced GABA decrease (measured in a separate session, inside the MRI scanner) correlated with the increase in AE 24 hours after receiving PA + left M1 a-tDCS (O’Shea et al. 2017), implicating this neurotransmitter in the mechanisms responsible for the memory enhancement. Therefore, we hypothesised that the solution to the temporal credit assignment might be reflected in the level of M1 excitation:inhibition during PA, such that greater E:I ratio would be associated with longer memory timescales. We used a weighted two-state model to infer this computation (i.e. relative contribution of short-lasting versus long-lasting computational states to the overall representation of the perturbation) at the individual level, as adaptation proceeded after or during real or sham left M1 a-tDCS. Individual trait markers of neurochemical response to M1 a-tDCS, white matter microsctructure, resting functional connectivity, as well as BDNF val66met polymorphism were taken to test the main prediction and explain individual variability in computation and behaviour. The data presented in this paper demonstrate that sensorimotor cortex E:I (and in particular GABAergic inhibition) causally determines the time constant of memory persistence, explaining phenotypic variation in adaptation decay in young healthy adults.

## Results

### Computational model of sensorimotor memory formation

To formally test the hypothesis that reduced M1 inhibition biases temporal credit assignment towards longer timescales, right-handed healthy volunteers (n = 24, mean age = 25 years, SD = 3.50) adapted to 10° right-shifting prisms, during or after real or sham left M1 anodal tDCS (4 sessions per individual, Figure 6). The PA paradigm was as before (O’Shea et al. 2017). Participants adapted gradually to prisms over 20 minutes (7 blocks, 120 trials, Figure 1B: E1-7). Leftward AEs were probed in interleaved blocks during PA (7 blocks, 15 trials each, Figure 1B: AE1-7). AE retention was assessed after a 10-minute rest delay (45 trials; Figure 1B: AE8-10). Unlike in most adaptation paradigms, in this PA protocol, error-dependent learning and AE are not quantified under near-identical conditions. Instead, two different tasks are used (Figure 1A, Figure 6). Learning is quantified, under prism exposure with visual feedback (Figure 1B: E1-E7), as endpoint accuracy on trials of speeded centre-out reaches to lateral targets. AE is measured, after prism removal without visual feedback (Figure 1B: AE1-7), as endpoint accuracy under non-speeded conditions to an untrained central target. By quantifying that portion of learning that transfers across a contextual change (of limb dynamics, speed, feedback, and spatial location), this AE measure reflects generalisation (Kitazawa et al. 1997), a factor thought to be an essential precursor of the cognitive transfer gains from PA observed in neurological patients with visual neglect (Jacquin-Courtois et al. 2013). Consistent with this, sensorimotor cortex stimulation stabilised both AE and cognitive gains in patients (O’Shea et al. 2017). Understanding the mechanisms of AE stabilisation would advance both motor neuroscience and therapeutics.

**Figure 1:**
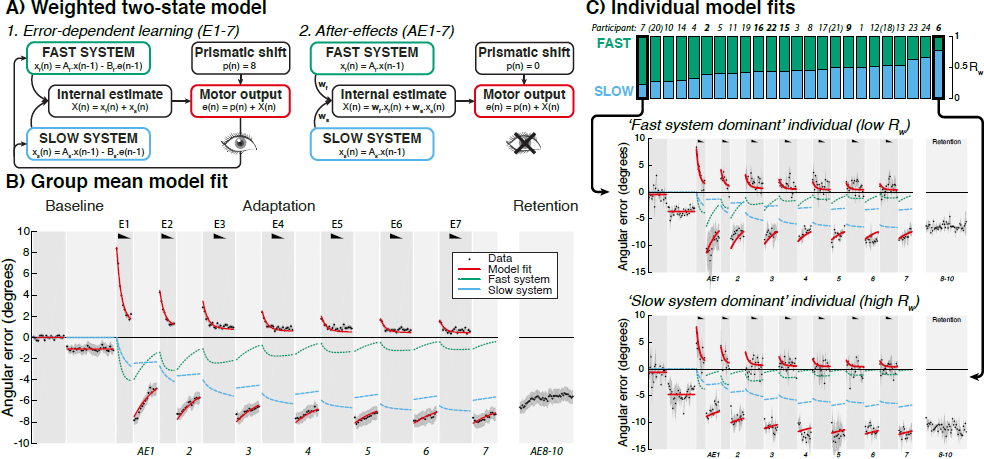
Computational modelling of motor behaviour during prism adaptation. **A.** Schematic representation of the weighted two-state model. **B.** Fit of the group mean prism adaptation data. The x-axis represents trial number and the y-axis pointing accuracy (rightward errors are positive). Black wedges indicate blocks throughout which prisms were worn. Group mean data (4 sessions per participants, i.e. 96 datasets) is plotted in black (error bar = SEM). The fitted weighted two-state model is represented in red and the time courses of the fast and slow systems in green and blue respectively. **C.** Individual fits in the absence of stimulation. Each bar plot represents the relative weight (*R_w_*) of one participant (estimated on the sham-tDCS conditions only), ranked in ascending order from left to right. Participant numbers are indicated on top of each bar (bold = Val/Met carriers, regular = Val/Val carriers, parenthesis = not genotyped). Individual model fits are plotted for the two extreme participants on this spectrum (lowest *R_w_* is S7; highest *R_w_* = S8). The individual with with lowest *R_w_* shows temporal dynamics of AE that more strongly resemble the fast (green) than the slow (blue) system, with steep slopes indicating a labile AE during Adaptation, which declines in magnitude across Adaptation to Retention. In the individual with highest *R_w_*, AE1-7 more strongly reflects the dynamics of the slow system (blue) than the fast (green), with flat slopes indicating stable AE during Adaptation, which increases in magnitude across Adapation to Retention.

To achieve this, we built a variant of an influential state-space model of adaptation that posits two adaptive processes learning from performance error – one ‘fast’ that learns and forgets quickly and one ‘slow’ that learns and forgets slowly Smith et al. 2006a. The net adaptation (during error-dependent learning and AE measurements) is the summed output of the two systems. During PA, error-dependent learning behaviour is relatively invariant across participants (Figure 1B: E1-7), however individuals differ in how the consequent AE develops and decays (Figure 1BC: AE1-7). The original model cannot account for this variability because the dynamics of AE behaviour are dependent fully upon the dynamics of error correction. To remedy this, we modelled the AE as a weighted sum of the two systems (Figure 1). Figure 1B shows this model provided a good fit to the group mean data (R^2^ = 99.68%), outperforming the original model after correcting for the benefit of two extra degrees of freedom (Δ*AIC* = 273.02; full details in Materials and Methods). When fitting individuals’ data, the learning and retention rates of both systems were kept constant for all. This approach reduced the number of free parameters, enabling individuals’ AE to be modelled solely in terms of the two weights, one per system. Within this modelling framework, temporal credit assignment can be conceptualised as the relative weight assigned to the slow versus fast system, computed as: *R_w_* = *w_s_*=(*w_s_* + *w_f_*)). Higher *R_w_* predicts more stable AE.

### Neuro-computational correlates of memory persistence

The model quantitatively captures inter-individual variation in PA profile, based on a principled theory of how (hidden) computational states contribute to (overt) motor behaviour. This suggests its parameters may be sensitive to the underlying underlying neural causes of this variation. To test this, we regressed the model parameters (*w_f_*, *w_s_*) against whole brain imaging measures of white matter microstructure and resting state functional connectivity. Figure 2 shows the structural and functional correlates of *w_s_* across participants (*w_f_* regressed out). Inter-individual variation in *w_s_* correlated with the microstructural integrity of the left superior longitudinal fasciculus (predominantly branches II and III), a white matter pathway interconnecting frontoparietal cortex (Figure 2A; *p* < 0.05, FWE-corrected with 2D-cluster enhancement) (Thiebaut De Schotten et al. 2011). The higher the weight an individual assigns to the slow system on AE, the stronger their coherence of white matter connectivity within this tract (i.e. higher fractional anisotropy and lower mean diffusivity).

**Figure 2:**
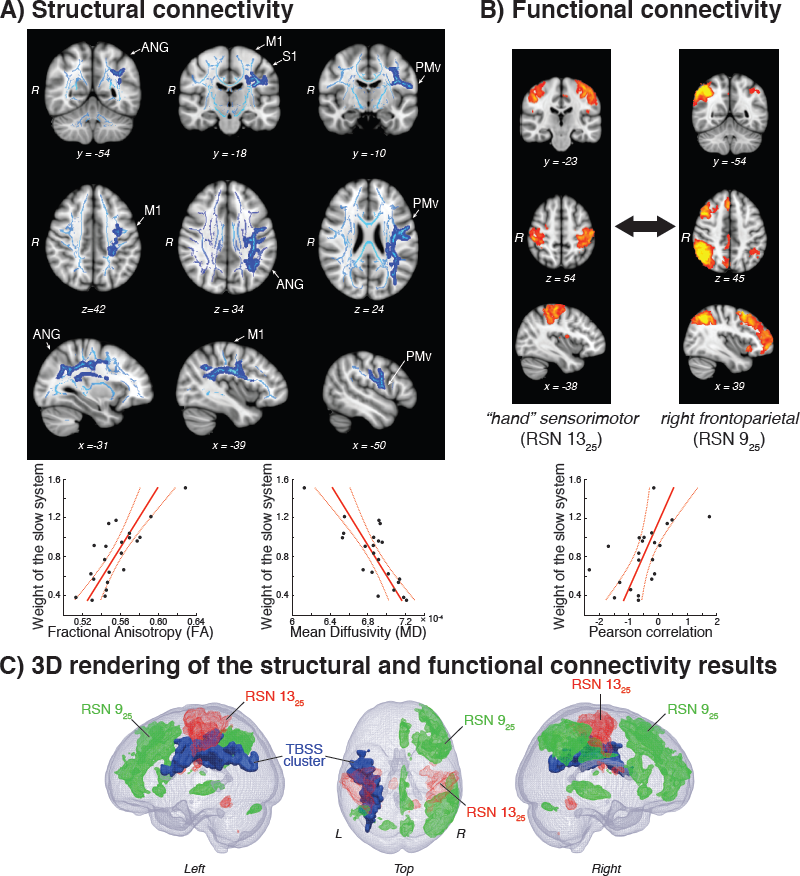
Structural and functional connectivity correlates of memory persistence. **A.** White matter whose microstructure covariates with the weight of the slow system (*w_s_*). Light blue shows the white matter FA skeleton in which statistical analysis (TBSS) was carried out. Dark blue indicates regions where FA is positively associated with *w_s_* and MD is negatively associated with *w_s_* (Fisher non-parametric combination, *p* < 0.05, FWE-corrected with 2D-cluster enhancement, *w_f_* regressed out). Liinear relationships are plotted underneath for illustration purpose. **B.** In the absence of stimulation, the degree of functional coupling between these 2 resting state networks (left: “hand” sensorimotor network, *Z* > 4, x = −38, y = −23, z = 54; right: right frontoparietal network, *Z* > 4, x = 39, y = −54, z = 45) was positively associated with the weight of the slow system (*p* < 0.05, FWE-corrected, *w_f_* regressed out). The relationship is plotted underneath for illustration purpose. **C.** 3D rendering of the structural and functional connectivity correlates of the weight of the slow system. On this glass brain are overlaid: 1) the aforementioned white matter voxels (in blue) whose microstructure is associated with *w_s_*, and 2) the “hand” sensorimotor network (in red, *Z* > 4) and right frontoparietal network (in green, *Z* > 4) whose functional connectivity is associated with *w_s_*.

Analyses of resting state data asked whether, across individuals, there was any functional network for which the strength of coupling with the “hand” sensorimotor network (RSN 13_25_; Figure 8) covaried quantitatively with *w_s_* (while controlling for *w_f_*). Figure 2B shows that only one network out of 12 showed this pattern: the right frontoparietal attention network (RSN 9_25_; Figure 8). Individuals with higher weight assigned to the slow system on AE measures during PA had greater functional coupling between the sensorimotor network and the right frontoparietal attention network in the resting state (edge 13_25_-9_25_; *p* = 0.038, FWE-corrected). These structural and functional relationships with *w_s_* were independent, and remained significant when analyses tested for one while controlling for the other (both *p* < 0.03). This suggests that trait variation in resting state structural and functional connectivity of parietal-sensorimotor-premotor circuits influences the (long) time constant of sensorimotor memory and decay.

### Sensorimotor cortex excitation-inhibition ratio causally determines memory persistence

Next, we tested our key mechanistic hypothesis: that excitation:inhibition balance in sensorimotor cortex causally determines whether AE persists or decays by biasing temporal credit assignment towards longer timescales. Figure 3A shows our neuro-computational hypotheses and one-way directional predictions about the causal relationships between sensorimotor cortex neurochemistry, temporal credit assignment, and behaviour at baseline (i.e. in the absence of excitatory stimulation). All the statistical analyses reported in this section were one-tailed given the a priori directional hypotheses set out in Figure 3A.

**Figure 3:**
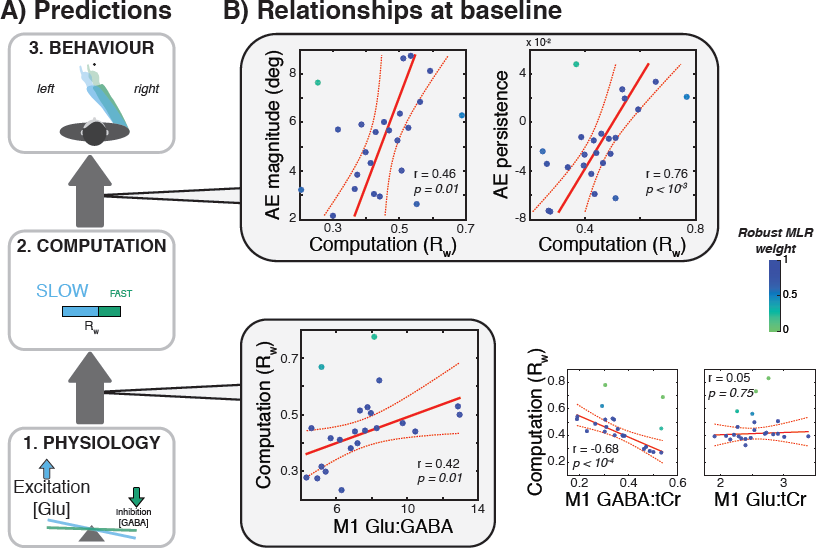
Predictions and correlations of MRS, computational and retention measures at baseline. **A.** Based on previous findings (Galea et al. 2010; O’Shea et al. 2017), we predicted greater M1 E:I ratio to be associated with greater relative contribution of the slow system to the AE during Adaptation, which in return should predict greater and more persistent AEs at retention. Analysis of baseline (i.e. in the absence of stimulation) inter-individual covariation in these 3 levels confirmed the predictions. Across individuals, higher Glu:GABA was associated with greater *R_w_* (*r* = 0.42; *p* = 0.01). Individual contribution of GABA and Glu to this relationship is plotted on the side. In return, greater *R_w_* during PA was associated with greater (*r* = 0.46; *p* = 0.01) and more persistent (*r* = 0.76; *p* < 10^−3^) AEs after a 10 minutes break. Colour code represents the contribution of individual data points to the linear regression.

First, we tested for a baseline relationship between sensorimotor cortex E:I and temporal credit assignment. Excitation:inhibition ratio was quantified across individuals as the relative concentration of glutamate to GABA in left sensorimotor cortex, using highfield (7T) resting state MRS. Across individuals, we predicted a positive correlation between E:I and the relative weight parameter (*R_w_*). Robust linear regression confirmed this (Glu:GABA *R_w_*: *r* = 0.42; *p* = 0.01; Figure 3B). When evaluating the individual contribution of each neurotrans-mitter to this relationship (analyses controlled for the other metabolite), greater GABA levels were found to be associated with lower *R_w_* (*r* = 0.68; *p* < 10^−4^). By contrast, glutamate showed no association with *R_w_* (*r* = 0.05; *p* = 0.75). In turn, as expected, higher *R_w_* during PA predicted greater and more persistent AE at retention (AE magnitude: *r* = 0.46; *p* = 0.01;AE persistent: *r* = 0.76; *p* < 10^−3^; Figure 3B).

To test the hypothesis that *R_w_* depends *causally* on E:I, we intervened within sensorimotor cortex with excitatory stimulation (a-tDCS) during PA. This has previously been shown to reduce GABA concentration (Stagg et al. 2009; Bachtiar et al. 2015; Antonenko et al. 2017). If E:I causally determines *R_w_*, then stimulation during PA should increase *R_w_* and consequently increase AE at Retention (Figure 4A). If the computation of *R_w_* is critical for this effect to arise, the same should not be observed when stimulation is delivered before PA (offline tDCS condition). In other words, across individuals, the magnitude of stimulation-induced neurochemical change (Δ E:I) should predict the computational change (Δ*R_w_*) induced by online (but not offline) a-tDCS, which should in return predict the behavioural change at retention (Δ*AE*).

**Figure 4:**
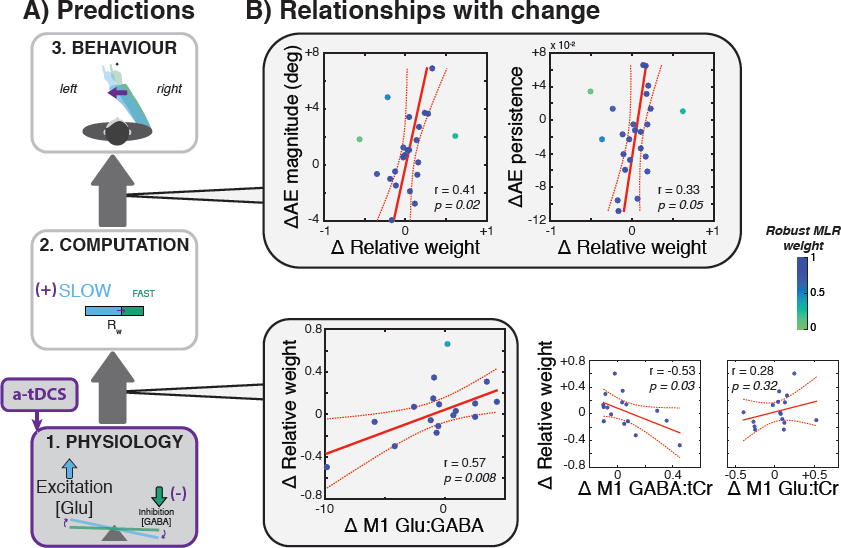
Predictions and correlations of stimulation-induced change in MRS, computational and retention measures. **A.** If the relationship described in Figure 3 are causal, then across individuals, the magnitude of stimulation-induced increase in E:I ratio should relate to how much *R_w_* increase, which should predict subsequent increase in AE at retention. **B.** Regression analyses confirmed these predictions (ΔGABA:Glu × *R_w_*: *r* = 0.57; *p* = 0.008; Δ AE magnitude: *r* = 0.41; *p* = 0.02; Δ AE persistence: *r* = 0.33; *p* = 0.05). Colour code represents the contribution of individual data points to the linear regression.

At the group mean level, stimulation did not change GABA concentration (*p* = 0.31; Figure 10), but stimulation during PA did increase AE at retention (*p* = 0.02, Cohen’s d = 0.4640; Figure 9A), while stimulation prior to PA did not (*p* = 0.64; Figure 9A), all three findings replicating previous work (full details in Appendix; O’Shea et al. 2017). Critically, analyses of inter-individual covariation in the effect of stimulation at the neurochemical, computational and behavioural level confirmed the predictions (Figure 4A). Across individuals, the greater the induced increase in Glu:GABA, the greater the increase in *R_w_* when a-tDCS was applied during PA (*r* = 0.57; *p* = 0.008, controlling for ΔGlu:GABA occurring with sham tDCS; Figure 4C). As expected, the stimulation-induced change in computation predicted behavioural change at retention. The greater the induced increase in *R_w_*, the greater the induced increase in AE at Retention (Δ AE magnitude: *r* = 0.41; *p* = 0.02; Δ AE persistence: *r* = 0.33; *p* = 0.05; Figure 4).

The analysis of the individual relationships of ΔGABA and ΔGlu with Δ*R_w_* (analysis controlling for the a-tDCS-induced change in the other metabolite and sham tDCS-induced change in Glu:GABA) revealed the same pattern found at baseline. Stimulation-induced decrease in GABAergic tone was associated with increase in *R_w_* (*r* = 0.53; *p* = 0.03) but changes in glutamatergic tone showed no such association (*r* = 0.28; *p* = 0.32). When stimulation was applied prior to PA, the a-tDCS-induced ΔE:I did not correlate with Δ*R_w_* (*r* = −0.20; *p* = 0.43). That induced ΔE:I increases AE retention, only when applied concurrent with (but not prior to) PA, implies causal dependence of stimulation on this computational state (*R_w_*). This confirms our mechanistic prediction: that higher sensorimotor cortex excitation biases temporal credit assignment towards longer timescales during PA, causing lasting memory.

### Enhanced memory persistence via sensorimotor cortex stimulation is modulated by genotype

Individuals with a common polymorphism in the gene coding for brain derived neurotrophic factor (BDNF val66met) exhibit reduced behavioural and neural markers of motor cortical plasticity (Kleim et al. 2006; McHughen et al. 2011; Joundi et al. 2012). Plastic enhancement of motor skill learning via a-tDCS is also reduced in Met allele carriers (Fritsch et al. 2010). The polymorphism causes a partial reduction in activity-dependent BDNF secretion (Egan et al. 2003; Chen et al. 2005), a factor involved in long-term potentiation (Minichiello 2009). Augmentation of BDNF-dependent synaptic plasticity is a candidate mechanism of action of sensorimotor cortex a-tDCS in mice and humans (Fritsch et al. 2010). Here we tested the prediction that this genotype would modulate the cognitive state dependent effect of stimulation on AE retention. That is, since the memory enhancement effect of sensorimotor cortex a-tDCS is activity-dependent (requires concurrent computation of *R_w_*), the impact of stimulation applied during PA should vary with genotype, while stimulation at rest prior to PA may not. Genotyping was acquired for 21/24 participants after all other data were analysed.

In agreement with the known allele distribution in the Caucasian population (Verhagen et al. 2010), 6/21 participants (28.6% of our sample) carried the Met allele. Mean AE at retention was analysed across all four PA sessions (anodal/sham tDCS × before/during PA). The interaction of genotype with cognitive state and stimulation was marginally significant (*F* (1; 19) = 4.3; *p* = 0.053, 2-tail, 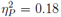; Figure 5). Follow-up ANOVAs confirmed that BDNF polymorphism influenced the behavioural response to stimulation when applied during PA (stimulation × genotype: *F* (1; 19) = 9.24; *p* < 0.01, 2-tail, 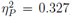) but not when applied prior to PA (*F* (1; 19) = 0.04; *p* = 0.84, 2-tail). As predicted, stimulation during PA significantly enhanced AE retention in Val-Val carriers (anodal-sham: *t*_(14)_ = 3.51; *p* < 0.01) but not in Val-Met carriers (*t*_(5)_ = 1.30; *p* = 0.25). These findings implicate activity-dependent BDNF secretion in stimulation-induced enhancement of AE persistence. They support the hypothesis that augmentation of BDNF-dependent synaptic plasticity is a contributory mechanism mediating behavioural plasticity induction via sensorimotor cortex anodal tDCS. Prior evidence was confined to motor skill learning. Our results extend this to sensorimotor adaptation.

**Figure 5:**
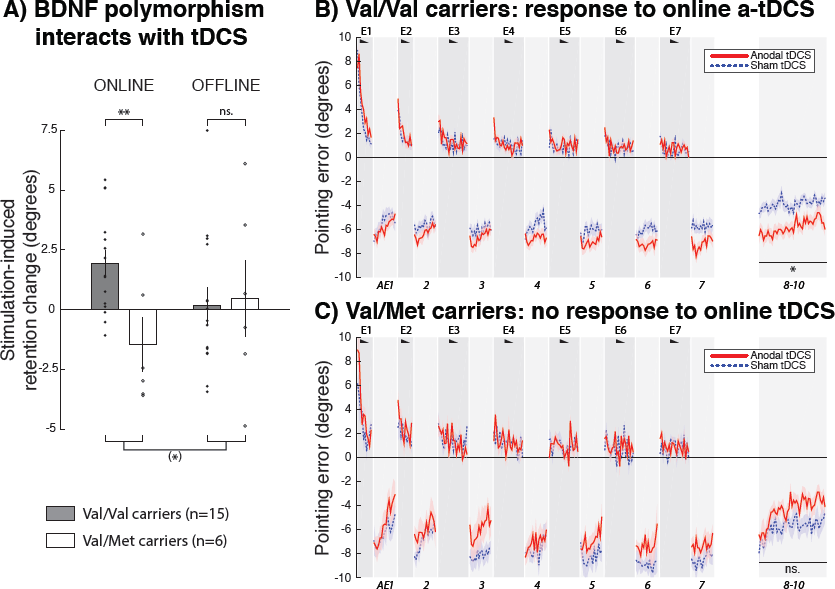
The BDNF Val66Met polymorphism influences the effect of anodal tDCS on retention. **A.** The change in prism AE magnitude at retention between anodal and sham tDCS (a value > 0 means an increased in AE) is plotted as a function of experiment (online/offline) and BDNF polymorphism (Val/Val in grey, Val/Met in white). Each dot represents an individual and the error bars represent the SEM. BDNF polymorphism influenced the a-tDCS effect only when stimulation was delivered during but not before PA. Val/Val carriers (n = 15) but not Val/Met carriers showed the expected increase in AE with online a-tDCS. Asterisk indicates significant effects ((*): *p* = 0.053; **: *p* < 0.01). **B-C.** Val/Val carriers but not Val/Met carriers responded to online M1 a-tDCS. The x-axis represents trial number and the y-axis pointing accuracy in reference to baseline accuracy (i.e. change from baseline, positive values indicate rightward errors). Pointing accuracy in Val/Val carriers (B, n = 15) and Val/Met carriers (C, n = 6) is plotted when anodal (red) or sham (blue) was applied to the left M1 during prism adaptation (online experiment) (group mean 1 SEM). Black wedges indicate blocks throughout which prisms were worn (CLP trials).

## Discussion

Adaptation is a fundamental property of the nervous system that underwrites the maintenance of successful actions across the lifespan (Shadmehr et al. 2010). Computational theories posit that individuals adapt to perturbations (e.g. lateral displacement of the visual field by prisms) by forming an internal model of it, used to adjust motor output and restore accuracy (Smith et al. 2006a). Once acquired, what determines whether adaptation memory persists or decays? The present study investigated this question from a computational and neurochemical perspective. We recently showed (and replicated in the present study) that excitatory stimulation (a-tDCS) of sensorimotor cortex during but not before PA increases retention of the AE (O’Shea et al. 2017). This timing-dependent effect indicates that understanding the information processing implemented within sensorimotor cortex and interacting with the stimulation is key to understanding the mechanism by which sensorimotor memories are formed and enhanced. Here, we hypothesised that sensorimotor cortex excitation:inhibition during PA might reflect the solution to the temporal credit assignment problem, such that lower inhibition would bias estimates towards longer timescales, and therefore longer-lasting AEs. To formally test this hypothesis, we inferred hidden computational states from behaviour using a variant of an influential state-space model (Smith et al. 2006a) that allowed us to quantify the relative contribution of short versus long timescale to the internal representation of the perturbation (Figure 1). This weighted two-state model outperformed the original model and provided a good fit to group mean and individual data. Across individuals, greater E:I ratio (quantified as Glutamate:GABA concentration using 7T MRS) was associated with greater contribution of the slow system during PA, which in return predicted greater and more persistent AEs at retention. To establish causality, the effect of excitatory a-tDCS was assessed on all three levels (physiology, computation, behaviour). This resulted in correlated increases in E:I, relative contribution of the slow system, and retention.

MR spectroscopy offers a way to measure metabolite concentrations in the living brain. The non-invasiveness of this technique comes at the cost of a relatively low signal-to-noise (SNR) ratio. In this study, good SNR was obtained by using ultra high-field (7T) MRS and by acquiring data from a large 2 × 2 × 2 cm^3^ MRS voxel centred on the region of interest. Due to the size of the voxel, adjacent regions of sensory cortex (S1) were also included in the measure of M1 metabolites. This is a common methodological limitation of MRS studies (Stagg et al. 2009; Lunghi et al. 2015; Antonenko et al. 2017; Ip et al. 2017; Kolasinski et al. 2017; O’Shea et al. 2017). Although we cannot rule out the contribution of S1 to our results, M1 is likely to play a predominant role because of a convergence of studies implicating it in the consolidation of adaptation memory (Wise et al. 1998; Li et al. 2001; Paz et al. 2003; Paz et al. 2005; Hadipour-Niktarash et al. 2007; Landi et al. 2011; Leow et al. 2016). It is not our intention however to suggest that only M1 is involved in the implementation of temporal credit assignment. As illustrated in the structural and functional connectivity results, the weight assigned to the slow system appears to involve the interaction of M1 with at least the posterior parietal cortex and ventral premotor cortex, two regions known to be involved in PA (Kurata and Hoshi 1999; Clower and Boussaoud 2000; Pisella et al. 2004; Newport and Jackson 2006; Newport et al. 2006; Danckert et al. 2008; Luauté et al. 2009). A parsimonious interpretation of the results is therefore that the level of inhibition within M1 is a causal factor in the computation of the timescale over which to retain the solution of the adaptation, but that it may also involve other distant regions.

The relationships between M1 inhibition and *R_w_* could result from two functionally distinct forms of GABAergic signalling: phasic or tonic. Phasic GABAergic activity refers to the release of presynaptic GABA into the synaptic cleft due to the depolarisation of inhibitory neurons (Mody 2001; Bachtiar and Stagg 2014). It is involved in the generation of rapid inhibitory postsynaptic currents (IPSCs) via the activation of GABA_B_ receptors (Maffei et al. 2017). In addition, extracellular GABA also has a longer-lasting neuromodulatory role via tonic signalling on extra-synaptic GABA_A_ receptors (Glykys and Mody 2007). This form of tonic inhibition affects neuronal excitability via an action on the tonic conductance, which is distinct from the time-dependent phasic inhibitory signalling (Mody 2001). It has an important regulatory influence on synaptic plasticity in that greater tonic inhibition is typically associated with less long-term potentiation (LTP) (Wigström and Gustafsson 1983; Chapman et al. 1998; Levkovitz et al. 1999; Bütefisch et al. 2000; Yoshiike et al. 2008; Martin et al. 2010). Extrasynaptic tonic GABA represents approximately 70% of the total GABA in the cortex (Petroff 2002) and, unlike vesicular phasic GABA, it is not bound to macromollecules, which makes it more easily detectable with MRS (Stagg et al. 2011b). Although these observations indicate that the resting MRS-GABA signal measured in this study reflects predominantly tonic extrasynaptic GABA_A_ inhibition (Stagg et al. 2011a; Stagg et al. 2011b), it is currently nearly impossible to disentangle the contribution of the two neurotransmitter pools with the resolution of this technique. Studies considering TMS intra-cortical inhibitory measures as indirect proxi of GABA_A_ and GABA_B_ inhibition have reported modulatory effects of a-tDCS on both types of signalling (Hummel et al. 2005; Antal et al. 2010; Tremblay et al. 2013; Amadi et al. 2015). We therefore suggest that the baseline relationship between ΔMRS-GABA and Δ*R_w_* is likely to rely on individual trait variation in GABA_A_ tone, while the relationship between ΔMRS-GABA and *R_w_* is likely to implicate state variation in both GABA_A_ and GABA_B_. The latter might contribute to explaining why although M1 a-tDCS has been shown to reduce MRS-GABA measures beyond the period of the stimulation (for up to 30 minutes) (Bachtiar et al. 2015), it had no effect on behaviour when applied before PA (O’Shea et al. 2017). Another account for the timing-dependent interaction between a-tDCS and sensorimotor memory is that offline a-tDCS might involve metaplasticity (Stagg et al. 2011c), a set of mechanisms engaged to counteract the effect of excitatory stimulation in order to maintain neural activity within a normal range (Lang et al. 2004). These mechanisms might take place during PA in the offline tDCS condition and suppress the excitatory effect of a-tDCS.

Identification of individual predictors of responsiveness to stimulation is crucial for both the mechanistic understanding of the effect of tDCS and the tailoring of interventions on an individual basis. In the present study, the cognitive state dependency of the stimulation-induced memory enhancement was influenced by individuals’ BDNF val66met polymorphism. Homozygote individuals (Val/Val) responded to online but not offline a-tDCS while heterozygotes individuals (Val/Met) responded to neither online nor offline a-tDCS. This result extends the known role of activity-dependent BDNF secretion in motor skill learning and its modulation by a-tDCS (Fritsch et al. 2010) to another paradigm: sensorimotor learning. It is unclear however how BDNF polymorphism interacts with the neuro-computational mechanism reported above. For example, it might be the case that because of altered activity-dependent synaptic plasticity, Met carriers show a different relationship between neurochemical and computational changes (e.g. a greater reduction in GABA is needed to induce the same increase in *R_w_* in Met carriers). The small number of Met carriers in our experimental sample (n = 6) prevented us from formally asking this question. Future research beyond the exploratory analysis reported here should therefore consider screening participants for their BDNF polymorphism at inclusion in order to obtain balanced sample sizes between the two groups (Val/Val and Val/Met) and formally compare the mechanisms involved.

## Materials and methods

### Participants

Twenty-four right-handed healthy male individuals (mean age = 25.03 years, SD = 3.50) gave their written informed consent to take part in this study in accordance with ethical approval from the Oxford A Research Ethics Committee (REC reference number: 13/SC/0163). The gender selection criterion was applied to avoid potential confounds related to variations in neurotransmitter concentration with the menstrual cycle in women (Smith et al. 1999; Epperson et al. 2002). Handedness was assessed using the Edinburgh handedness questionnaire (Oldfield 1971). All participants had normal or corrected-to-normal vision and indicated no family history of psychiatric or neurological disease. All participants were naive to the purpose of the experiment and were not informed of the expected effect of the prism glasses and brain stimulation. No participant reported any side effect from the experimental procedure.

### Experimental design

All individuals participated in four PA sessions (behavioural experiment) and two scanning sessions (neuroimaging experiment). Sessions were separated by at least one week. The order of the four PA sessions was fully counterbalanced across the group. The order of the scanning sessions was also counterbalanced across the group.

The behavioural experiment had a 2-by-2 within-subject design. Experimental sessions differed in regards to the type (anodal or sham tDCS) and timing (during or before PA, i.e. online or offline) of the stimulation. Similar to our previous report (O’Shea et al. 2017), each PA session included 4 phases as follow: 1) measure of baseline pointing accuracy (3 blocks); 2) prism adaptation (14 blocks; 20 min); 4) 10-minute break (blindfolded, at rest); 3) measure of prism AE retention (Figure 6B). The neuroimaging experiment (performed in the same individuals) involved 2 scanning sessions with either real or sham left M1 tDCS. Details of the scanning protocol is provided in Figure 6C.

**Figure 6:**
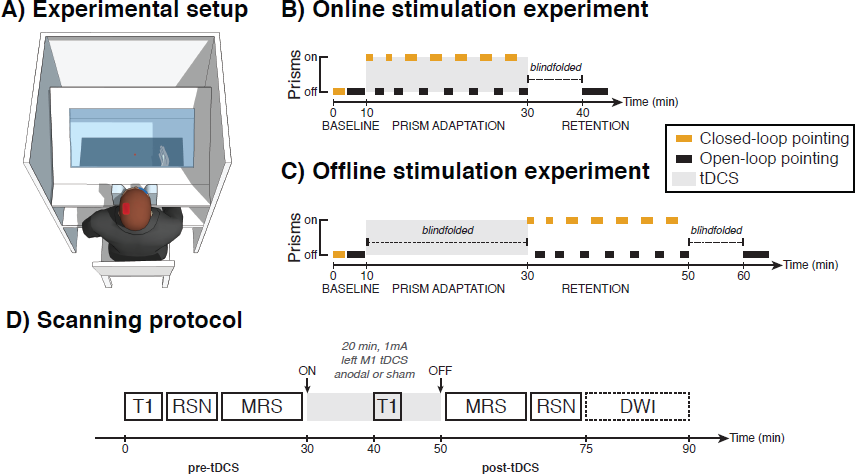
Experimental set-up and protocols. **A.** An PA set-up was used (see Materials and Methods). **B-C**) Experimental protocol. Closed-loop pointing (CLP) and open-loop pointing (OLP) accuracy was first measured at baseline. Then followed 20 minutes of either real or sham left M1 a-tDCS during which participants either adapted to 10-degree right-shifting prisms (online condition, B) or stayed blindfolded at rest (offline condition, C). In the offline condition, participants underwent prism adaptation as soon as the stimulation finished. The prism adaptation phase included blocks of CLP during which participants adapted to prisms, interleaved with blocks of OLP used to measure the development of prism AE. After the end of the last block of the PA phase, participants rested blindfolded for 10 minutes before to undergo a long block of 45 OLP trials to measure retention. **D.** Scanning protocol. All participants underwent two scanning sessions with either anodal or sham left M1tDCS, the order of which was counterbalanced across the group. MR spectroscopy and resting state fMRI were acquired at two time points: before and after tDCS. Diffusion-weighted images were taken once at the end of one of the two scanning sessions in 22 subjects. Timeline shown is an estimate of the length of the scans in minutes.

### Prism adaptation procedure

The experimental protocol used in this study was as described in O’Shea et al. (2017) and is summarised bellow. The main difference with previous work (O’Shea et al. 2017) is that a fully automated experimental set-up was used to enable a more precise and reproducible measurement of reaching errors, as well as a better control of visual feedback availability and experimental timing (Figure 6A). As a result, the pace of the experiment increased so we could fit 7 exposure blocks instead of the original 6 during the 20 minutes of brain stimulation (Figure 6BC).

Participants sat facing a horizontal 32-inch LED screen embedded in a table that was used to record the reach endpoint position of their index finger (Figure 6A). Their head was restrained by a chin rest mounted on the edge of the table to keep a distance of about 60 centimetres between their eyes and the centre of the screen. Wood panels were placed on the other three edges of the table to create a box in which participants’ visual environment was strictly restricted to the screen. A button was attached to the pole of the chin rest and subjects were instructed to keep it pressed at all time and only release it when initiating a reaching movement. A fixed shutter prevented participants from seeing their limb when pressing the button and during the first third of their pointing movement. A liquid crystal shutter (Dispersion film, Liquid Crystal Technologies, Ohio, USA) placed between participants’ eyes and their limb controlled the visual feedback of the screen and pointing limb. The same prism glasses were used (glacier goggles: Julbo, Longchaumois, France; 10° right-shifting prism lenses: OptiquePeter, Lyon, France).

Participants were instructed to perform two types of pointing movements: closed-loop pointing and open-loop pointing.

- **Closed-loop Pointing (LP).** On CLP trials, participants made speeded reaching movements (average movement duration = 317 ms, SD = 53 ms) to point at a visual target appearing on the screen at the beginning of each trial. Within any CLP block, targets appeared either 12 cm to the left or 12 cm to the right of the centre of the screen (50% on each side), following a pseudorandom sequence in which target location transitions were generated if the same location appeared twice in a row. Visual feedback was limited to the last two thirds of the reaching movement in order to limit strategic adjustment (Redding and Wallace 1996; O’Shea et al. 2014; Inoue et al. 2015; O’Shea et al. 2017) and lasted until 500 ms after participants touched the screen to allow for them to perceive their terminal reach endpoint error. After this time, the LC shutter turned opaque and participants had to return to the starting position (i.e. press and hold the button) without visual feedback of their hand.
- **Open-loop pointing (OLP).** On OLP trials, participants pointed at a slower speed (average movement duration = 816 ms, SD = 174 ms) to a visual target located in the centre of the screen. The LC shutter turned opaque as soon as their hand left the starting position (i.e. release of the button), thereby blocking any visual feedback during the reaching and return movements. The LC shutter turned clear again only when participants returned to the starting position and initiated the next trial.

Four factors differentiated OLP from CLP trials: 1) participants only wore prism glasses during CLP trials (and never during OLP trials) so that any change in pointing accuracy on open-loop pointing relative to baseline could be interpreted as an after-effect of the adaptation; 2 participants could learn from their reach endpoint error on CLP but not on OLP (because visual feedback was deprived); 3) participants made fast movements on CLP (average movement duration = 317 ms, SD = 53 ms) but slower movements on OLP (average movement duration = 816 ms, SD = 174 ms); 4) participants pointed to lateral targets on CLP (located 12 cm to the left or right of the centre of the screen) and to a unique central target on OLP. The differences in movement speed and target location between prism exposure (CLP trials) and AE probes (OLP) enabled us to assay generalised AE that were not contaminated by extensive training of a specific movement dynamic and local learning at a certain target location (Kitazawa et al. 1997). This type of AE is thought to be relevant for neglect rehabilitation (Serino et al. 2006; O’Shea et al. 2017).

In all four PA sessions, the tDCS electrodes were positioned and the stimulation switched on at the end of the baseline phase (20 CLPs, 30 OLPs). In the online condition, the PA phase (E1-7: 10 CLPs per block during the first 2 blocks, 20 CLPs per block in the remaining ones; AE1-7: 15 OLPs per block) started as soon as the stimulation reached its constant level. In the offline condition, participants were blindfolded and remained at rest during the entire duration of the stimulation. This enabled us to control the visual input participants received during the offline stimulation. The prism adaptation phase started as soon as the stimulation was over in the offline condition. In all sessions, retention of the AE (AE8-10: 15 trials per block) was probed 10 minutes after the end of the last OLP trial of the adaptation phase.

### Transcranial direct current stimulation

#### During behavioural sessions

When delivered outside the scanner (i.e. PA sessions), direct current stimulation was generated by a battery driven DC stimulator (Neuroconn GmbH, Ilmenau, Germany) connected to two 7 × 5 cm sponge electrodes soaked in a 0.9% saline solution. The anode electrode was centred over C3 (5 cm lateral to Cz) corresponding to the left primary motor cortex according to the international 10-20 System (Herwig et al. 2003). The cathode electrode was placed over the right supraorbital ridge. The large surface of the electrode (35 cm2) ensured a good coverage of the M1 “hotspot”. Similar to previous M1 tDCS studies, the long axis of the anode electrode was oriented parallel to the sagittal axis during behavioural sessions (Nitsche and Paulus 2000; Galea et al. 2010; Panouillères et al. 2015; O’Shea et al. 2017). The electrodes were positioned immediately before stimulation onset and removed as soon at the stimulation finished in order to minimise participants’ discomfort. For anodal stimulation, the current intensity was set to 1 mA for 20 minutes with a ramp-up and ramp-down period of 10 seconds prior and after the stimulation respectively. For sham stimulation, the current also ramped up and down for 10 secs but no stimulation was delivered during the 20 minutes. Instead, small current pulses (110 µA over 15 ms) occurred every 550 ms to simulate the tingling sensation associated with real anodal stimulation. Both experimenters and participants were blinded to the stimulation condition. In order to verify the efficacy of the blinding, participants were asked to guess whether the stimulation was ‘real’/ or ‘fake’at the end of every behavioural session. The proportion of correct responses for identifying the stimulation condition was at chance level (50%, SD = 27.59), indicating that participants were unable to successfully differentiate anodal from sham tDCS.

#### Inside the scanner

When delivered outside the scanner (i.e. PA sessions), direct current stimulation was generated by a battery driven DC stimulator (Neuroconn GmbH, Ilmenau, Germany) connected to two 7 × 5 cm MR-compatible carbon-rubber electrodes equipped with an inbuilt 5 kΩ resistor at the junction between the electrode and the wire (Easycap GmbH, Herrsching, Germany) in order to avoid temperature increase and reduce induction voltages caused by the scanner (Woods et al. 2016). A thick coating of chloride-free abrasive electrolyte gel was applied between the biocarbon electrodes and the scalp to maintain low levels of impedance throughout the duration of the scan and ensure the stimulation to be delivered evenly across the electrode (Woods et al. 2016). The electrode placement was identical except for the anode electrode that was oriented perpendicular to the sagittal axis (instead of parallel, during behavioural sessions) in order for the wire to exit the RF coil through the shortest route and limit the risk of eddy current induction (Woods et al. 2016). This electrode montage was similar to previous work done in the laboratory (Stagg et al. 2009; Bachtiar et al. 2015; O’Shea et al. 2017). The electrodes were placed on participants’ head immediately before the beginning of the scanning session and removed as soon as participants exited the scanner. The protocol for anodal and sham stimulation was identical to what was done outside the scanner. Only participants were blinded to the stimulation condition inside the scanner.

### Statistical analysis of raw prism adaptation data

Reach endpoint angular errors were calculated as the angle formed between a straight line joining the starting position (button) and the target location and a straight line joining the starting position (button) and the touch location on the screen. Data distribution assumption of normality was assessed quantitatively using the Kolmogorov-Smirnov and Shapiro-Wilkinson tests and qualitatively using distribution plots and Q-Q plots. Sphericity was assessed using Mauchly’s test, with violations corrected using the Huyhn-Feldt procedure where appropriate. When data met the assumptions of parametric testing, pointing angular error data were analysed with repeated measures ANOVA and t-tests using SPSS software (SPSS inc. version 24.0) and Matlab (The MathWorks inc. version R2014b). Cohen’s d was reported for significant results only. All the MATLAB scripts are provided in supplementary material.

### Computational analysis of prism adaptation data

State-space models and Bayesian models have proven useful in describing and predicting motor behaviour during sensorimotor adaptation (Kitazawa et al. 1995; Körding and Wolpert 2004; Smith et al. 2006a; Körding et al. 2007; Berniker and Kording 2008; Joiner and Smith 2008; Haith et al. 2009; Lee and Schweighofer 2009; Yamamoto and Ando 2012; Inoue et al. 2015; McDougle et al. 2015). One key difference between these two classes of model is that the former extracts hidden internal states from the raw behavioural data whereas the latter infers them from the noise structure of the experiment under the assumption of optimality. Bayesian models therefore require the reliability of different sources of information (e.g. vision, proprioception) to be experimentally manipulated (for examples of such experimental manipulations, see Körding and Wolpert 2004; Wei and Körding 2010). The set of experiments reported in this thesis was designed to measure the evolving dynamics of the AE during and after adaptation to prisms and did not involve manipulations of the reliability of different sources of information. Hence the state-space modelling framework was used.

State-space models posit that trial-by-trial adaptation arises from the interaction of multiple learning processes that can be approximated by two systems: a ‘fast’ system that learns quickly but also forgets quickly, which is mainly responsible for the rapid initial error correction, and a ‘slow’ system that learns slowly but forget slower, allowing the long-term retention of AE (Smith et al. 2006a; Joiner and Smith 2008). A recent study applied time-invariant state-space models to prism adaptation data and suggested the need of a third ‘ultra-slow’ system when explaining the temporal dynamics of AE after prolonged prism exposure (500 exposure trials) (Inoue et al. 2015). The present experiment however used a relatively short prism exposure (120 pointing trials), which does not require the addition of a third system. On CLP, the two-state model is defined as follows:

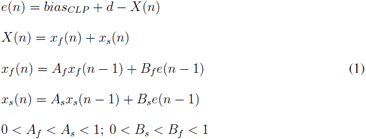

where the endpoint error on the *n*-th trial *e*(*n*) is modelled as the sum of a bias term *bias_CLP_* and a prism effect *d* from which is removed the sum of the states of the two systems *X*(*n*). The bias term is a fixed parameter capturing the natural deviation from target participants show at baseline (i.e. a constant ‘irreducible’ error that is not attributable to PA). The prism effect *d* is a fixed parameter that determines the error magnitude obtained on the first prism exposure CLP trial of the Adaptation. Owing to online corrections during reach trajectory with visual feedback (CLP), *d* is always less than the true visual displacement, and is typically about 60-80% (Rock et al. 1966; Redding and Wallace 2000; Redding et al. 2005). The total amount of adaptation *X*(*n*) corresponds to the sum of the states of the fast system *x_f_*(*n*) and slow system *x_s_*(*n*). These systems produce a trial-by-trial estimate of *d* based on the visual error feedback. The learning rate *B* dictates the proportion of error added to the state of a system from one trial to the other. A greater *B* means faster learning, i.e. larger proportion of error on the *(n-1)-th* trial corrected for on trial *n*. The retention rate *A* determines the speed of the memory decay, i.e. the proportion of state that is being carried over from one trial to the other. A high *A* for a system means little intertrial memory decay.

On OLP trials, where the visual disturbance is removed and no visual feedback is available, the original two-state model (Smith et al. 2006a) becomes:

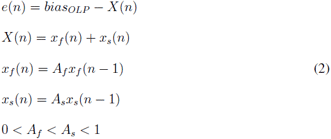

This equation differs from equation 1 in three ways. First, a different bias term is used, reflecting the fact that, at baseline, individuals show different biases between the two tasks (for example, compare the baseline deviation on the CLP and OLP blocks in Figure 1C). Second, the prism effect *d* is removed (effectively set to zero) because individuals are no longer wearing prisms when the AE is measured. Third, the learning rates of the fast and slow systems are set to zero (i.e. *B_f_* = 0; *B_s_* = 0) to reflect the absence of learning during OLP trials. Any within-block change in pointing accuracy during AE measurements is therefore explained by the forgetting of the two systems in this model. If visual feedback is provided during AE measurements (i.e. CLP, e.g. de-adaptation phase of experiment 1), the learning of both systems is switched on and the equation is the same as in 1 except that *d* = 0. In this case, within-block change in pointing accuracy reflects both the forgetting of the adapted state and the active learning from performance error (i.e. de-adaptation).

To date, state-space models have been used predominantly in the literature on visuomotor rotation (Lee and Schweighofer 2009; Tanaka et al. 2009; Tanaka et al. 2012; Kim et al. 2015; McDougle et al. 2015) and force-field adaptation experiments (Smith et al. 2006a; Joiner and Smith 2008), two paradigms in which the AEs are typically measured using the same experimental constraints as during exposure to the perturbation (i.e. identical effectors, movement dynamics, target location in space). Because of these identical conditions, both error-dependent learning and AE measurements are assumed to reflect the direct sum of the states of the two systems (see equations 1 and 2) (Smith et al. 2006a). However, if the error-dependent learning and AE measurements occur under different pointing conditions (CLP vs. OLP: different target location, different movement speed), the assumption of equal contribution of the two systems to the AE may no longer hold. For example, one system may generalise more than the other. In order to capture this key feature of our experimental paradigm in computational terms, we introduced a time-invariant weighting on the two systems that determines how much of what a system learns during prism exposure is subsequently expressed behaviourally on open loop pointing. In practice the equation for CLP trials remains the same as the original model (equation 1). However, the equation for OLP becomes:

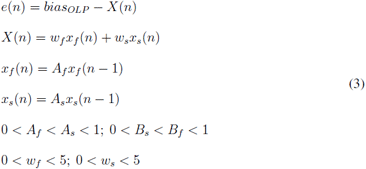

where *w_f_* and *w_s_* are weighting factors that introduce a dissociation between the state of a given system on CLP and its contribution to the total amount of adaptation on OLP. As a result, the AE is now modelled as a weighted sum of the states of the two systems. For each system, having a weight lower than 1 means a scaling down of its contribution to the prism AE, whereas a weight greater than 1 means an over-expression.

Model optimisation was performed in MATLAB (The MathWorks inc. version R2014b) using a least squares fitting procedure (*fmincon* function). In order to confirm the superiority of the weighted two-state model over the original non-weighted two-state model (Smith et al. 2006a), we performed formal model comparison on the group mean data (average of the 4 sessions of the 24 participants, i.e. 96 datasets). For the original two-state model, the best-fitted parameters were Af = 0.866; As = 0.991; Bf = 0.195 and Bs = 0.101 and for the weighted two-state model, they were Af = 0.931; As = 0.996; Bf = 0.159; Bs = 0.076; wf = 0.974 and ws = 0.855 (Supplementary scripts). The overall fit of the group mean data was better for the weighted two-state model compared to the original model (root mean squared erro*r* = 0.23° vs. 15.48°; R-square = 99.68% vs. 99.14%). In order to formally compare the two models, Akaike’s Information Criterion (AIC) and the associated relative likelihood (RL) were calculated for each model as follow:

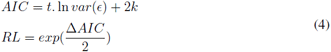

where t is the total number of data points and k is the degrees of freedom in each model (k = 4 for the original model and k = 6 for the weighted model). This index expresses the residual error while adding a penalty for model complexity in order to avoid issues related to over-fitting. Thus, the best model is characterised by the lowest AIC (i.e. a negative Δ*AIC*). This analysis confirmed that the improvement associated with the weighted two-state model was not merely due to extra parameters (Δ*AIC* = ‒273.02, *RL* = 3.06*x*10 ‒ 61).

The main advantage of the weighted two-state model is not only that it provides a closer fit to the data, but mainly that it enables to extract the key computational parameter of interest for this study, i.e. the time-invariant contribution of the fast and slow systems to overt behaviour on OLP during Adaptation. We expressed this as a relative weight as follow:

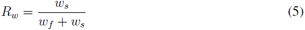

This derived scalar captures in a single metric the relative contribution of long versus short memory timescales to the AE during Adaptation. A relative weight greater than 0.5 is indicative of a dominance of the slow system, i.e. little intra- and inter- block forgetting (Figure 1C). Alternatively a fast system dominant PA profile (*R_w_* < 0.5) is characterised by larger prism AE initially but greater forgetting throughout the 20 minutes of prism adaptation (Figure 1C).

In order to ensure comparability of the relative weights within and between participants, *A_f_*, *B_f_*, *A_s_* and *B_s_* were kept constant and only *w_f_* and *w_s_* were estimated when fitting individual datasets. We used the learning and retention factors extracted from the fit of the weighted two-state model to the group mean data. If all parameters were let free, weighting factors would refer to different memory timescales depending on which participant/session they apply to. Instead, we decided to posit identical systems for all participants and only let the weight of these systems vary. This approach also reduced the number of free parameters, making the estimates more robust and ensuring that they capture unique variance. Furthermore, inspection of raw PA data revealed a stereotypical behavioural profile on CLP while most of the variance was present in the OLP data (Figure 1B).

### Regression analyses

Regression analyses were performed in MATLAB (The MathWorks inc. version R2014b) using robust multiple linear regressions analyses with an iteratively re-weighted least squares procedure. This algorithm was used to limit the contribution of potential outliers to the fit. Individuals with missing data for at least one of the variables included in the linear regression were excluded from the analysis. The details of participants exclusion is provided in Table 1. For all multiple linear regressions, normal probability plots of residuals were examined to check the normal distribution of errors. The coefficients reported are the regression coefficient estimates after z-transformation of the dependent and independent variables. Fisher’s r was calculated to compare regression coefficients.

**Table 1:**
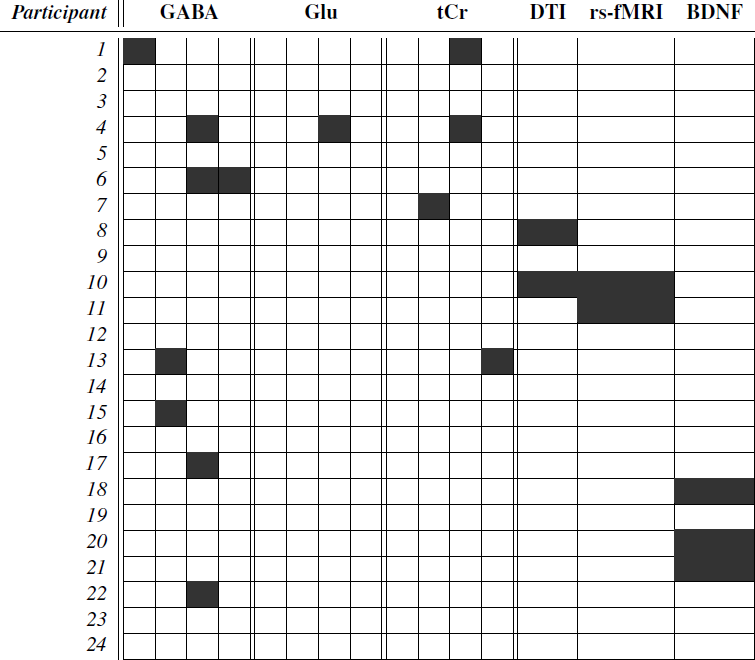
Diagram of data completeness. Completeness of the dataset is represented for each participant. The four columns of GABA, Glutamate and Creatine represent the four measurements (before a-tDCS, after a-tDCS, before s-tDCS, after s-tDCS). A black square means that the data didn’t meet the quality criteria. Black squares in the BDNF column indicates participants for which genotyping could not be obtained.

### BDNF Val66Met polymorphism

A sub-sample of 21 participants (Table 1) was genotyped for the BDNF Val66Met polymorphism (rs6265, G > A). Genomic DNA was extracted from buccal cells using the ChargeSwitch® gDNA Buccal Cell Kit (ThermoFisher Scientific, UK) and samples were genotyped in duplicate by LGC Genomics (LGC Group, UK). Rs6265 was the only polymorphism examined.

### MR data acquisition

All MR data (T1-weighted images, MR spectroscopy, diffusion-weighted images and resting state fMRI) were acquired on a 7T Siemens MAGNETOM scanner (Siemens, Erlangen, Germany) at the Oxford Centre for Functional Magnetic Resonance Imaging of the Brain (FMRIB). All participants underwent two 90 minutes scanning sessions (anodal tDCS scan and sham tDCS scan) separated by at least 1 week and counterbalanced across participants (Figure 6D). To keep volunteers engaged during the scanning session, a video (BBC life, no sound, no subtitles) was projected in the scanner at the exception of during the resting state fMRI acquisitions. Participants wore earplugs throughout scanning.

#### T1-weighted structural images

High-resolution T1 images (MPRAGE: 176 x 1 mm sagittal slices; TR/TE = 2200/2.82 ms; flip angle = 7 deg; FOV = 256 x 256; voxel size = 1mm isotropic; PAT = 2; scan time = 325 secs) were acquired at the beginning of each scanning session for MRS voxel placement and registration purposes.

#### MR spectroscopy

MR spectroscopy was acquired immediately before and after the application of 20 minutes of left M1 real or sham a-tDCS inside the scanner. B0 shimming was performed in a two-step process. First, GRE-SHIM (64 x 4mm axial slices, TR = 600 ms; TE1/2 = 2.04/4.08 ms; flip angle = 15 deg; FOV = 384 × 384; voxel size = 4 mm isotropic; scan time = 44 secs) was used to determine the optimal first- and second-order shim current. The second step involved the fine adjustment of first-order shims using FASTMAP. Prior to switching on the direct current stimulator, the modified semi-LASER sequence (O’Shea et al. 2007; Bank et al. 2015) (TR/TE = 5000/36 ms, 64 scan averages, scan time = 320 secs) was used to acquire MRS measurement in a 2 × 2 × 2 *cm*^3^ volume of interest centred on the left motor knob (Yousry et al., 1997) to include parts of the pre- and post-central gyrus (see VOI overlap maps in Figure 10B). As soon as the baseline MRS measurement was over, the experimenter turned on the DC stimulator and no scanning happened for 10 minutes (Figure 6D). At the end of this time, lower-resolution T1-weighted images (MPRAGE: 176 × 1 mm sagittal slices; TR/TE = 2200/2.82 ms; flip angle = 7 deg; FOV = 192 × 192; voxel size = 1 mm isotropic; PAT = 4; scan time = 171 secs) were acquired to check for head motion. If significant head motion was detected, the VOI was re-positioned to match the baseline acquisition and B0 shimming was performed again. If no significant head motion was detected, baseline VOI positioning and shimming were applied. On average, post-tDCS MRS measurements (64 averages) started 2 minutes 21 seconds after the end of the stimulation (range: 6 secs to 12 minutes 30 secs).

#### Diffusion weighted images (DWI)

Diffusion-weighted echo planar images (2 acquisitions of 64 diffusion-weighted directions in opposite phase-encode direction, b-value = 1000 s/mm2; 8 non-diffusion weighted images with 4 b0 volumes in the reverse phase-encode direction, b-value = 0 s/mm2; 120 slices; TR/TE = 6210/68.2 ms; FOV = 192 × 192; voxel size = 1.2 mm isotropic; PAT = 2; scan time = 938 secs) were acquired once at the end of a scanning session for 22 participants (Figure 6D; Table 1).

#### Resting state functional MRI (rs-fMRI)

Resting state Blood oxygen-level-dependent (BOLD) echo planar fMRI scans (96 slices; TR/TE = 1472/25.0 ms; FOV = 192 × 192; voxel size = 1.5 mm isotropic; PAT = 2; scan time = 444 secs) were acquired before and after the stimulation (Figure 6D). Participants were instructed to keep their eyes open and fixate on a cross hair projected on the screen. Immediately after each resting state fMRI scan, a field map (64 slices; TR = 620 ms; TE1/2 = 4.08/5.1 ms; FOV = 192 × 192; voxel size = 2 mm isotropic; scan time = 122 secs) was acquired for post-processing correction of geometrical distortions caused by magnetic field inhomogeneity. Two participants did not undergo the resting state scans in order to minimise the time they spent in the scanner (Table 1).

### MR data analysis

#### MR spectroscopy

Metabolites were quantified using LCModel (Provencher 1993; Provencher 2001; Provencher 2012) performed on all spectra within the chemical shift range 0.5 to 4.2 ppm. The model spectra of alanine (Ala), ascorbate/vitamin C (Asc), aspartate (Asp), glycerophosphocholine (GPC), phosphocholine (PCho), creatine (Cr), phosphocreatine (PCr), GABA, glucose (Glc), glutamine (Gln), glutamate (Glu), glutathione (GSH), myo-inositol (myo-Ins), Lactate, N-acetylaspartate (NAA), N-acetylaspartylglutamate (NAAG), phosphoethanolamine (PE), scyllo-inositol (scyllo-Ins) and taurine (Tau) were generated based on previously reported chemical shifts and coupling constants by VeSPA Project (Versatile Stimulation, Pulses and Analysis). The unsuppressed water signal acquired from the volume of interest was used to remove eddy current effects and to reconstruct the phased array spectra (Natt et al. 2005). Single scan spectra were corrected for frequency and phase variations induced by subject motion before summation. The full width half maximum (FWHM) and signal-to-noise ratio (SNR) did not differ between the four MRS measurements (Table 2).

**Table 2:**
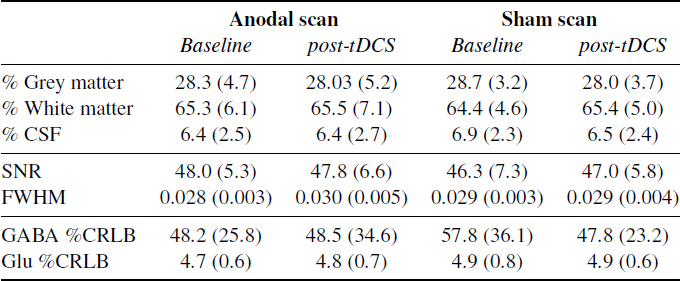
MRS characteristics. Group mean (standard deviation in parenthesis) voxel composition, spectrum signal-to-noise ratio (SNR), full width at half maximum (FWHM), and GABA and glutamate (Glu) relative Cramér-Rao lower bound (%CRLB) are shown for each spectroscopy measurement (before a-tDCS, after a-tDCS, before s-tDCS, after s-tDCS).

Relative Cramér-Rao Lower Bounds (CRLB) is commonly used as a quality-filtering criterion to identify and discard ‘bad quality’ data (Provencher 1993; Provencher 2001; Scholz et al. 2009; Kim et al. 2014; Antonenko et al. 2017). However, as recently highlighted in the work of Kreis (2016), this method induces a selection bias towards high concentration estimates and therefore leads to an over-estimation of the group mean metabolite concentration. This is particularly problematic for low concentration metabolites such as GABA. In order to avoid this methodological confound in this experiment, we used the alternative quality filtering method. First, a high relative CRLB threshold of 100% was applied in order to exclude exceptionally unreliable measurements. Next, we calculated the expected relative CRLB value for all MRS measurements under the assumption of a constant noise level across all measurements. We then excluded measurements for which the observed relative CRLB was exceptionally higher than the one predicted under this assumption. We used Pearson residuals to quantify the distance between the observed and predicted relative CRLB (threshold = 2). This quality filtering method allowed us to identify poor quality measurements irrespective to their concentration estimate and therefore avoid the selection bias of the standard method based on relative CRLB. Out of the 24 participants, 16 passed this quality check successfully for GABA, glutamate and creatine across all four MRS measurements. Twenty-three participants had at least one reliable baseline MRS measurement for all these metabolites (Table 1).

**Figure 7:**
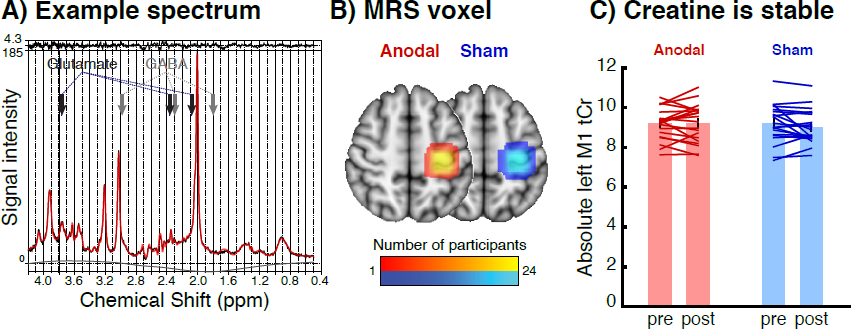
Magnetic Resonance Spectroscopy data. **A.** Example raw spectroscopy spectrum and LC model fit from one participant. The fitted LCModel (red) is overlaid on the raw data (black). The difference between the data and model (residuals) is shown at the top, and the baseline is shown at the bottom. The arrows represent the constituent peaks of glutamate (blue) and GABA (green). **B.** MRS voxel group overlap map. The MRS was centred on the left motor knob to include parts of the pre- and post-central gyrus (Yousry et al. 1997). This resulted in consistent positioning of the voxel in the same location in MNI space between individuals (light colours indicate overlap in voxel position in all 24 participants) and scanning sessions (red: anodal tDCS scan; blue: sham tDCS scan). **C.** The total creatine (tCr) concentration estimate was stable across all 4 measurements. Total creatine’s absence of significant variation across measurements (*F* (1; 19) = 1.46; *p* = 0.24) and good test-retest reliability (*ICC* = 0.86; *p* < 10^−3^) enabled us to use it for internal referencing.

FMRIB’s automated segmentation tool (Patenaude et al. 2011), part of the FSL software library (Smith et al. 2004), was used on the T1-weighted images to calculate the percentage of grey matter, white matter and cerebrospinal fluid present in the volume of interest (noted [GM], [WM] and [CSF] respectively) (Table 2). GABA and glutamate concentration estimates were corrected for the proportion of grey matter within the VOI (i.e. divided by [GM]/([GM] + [WM] + [CSF])), and creatine and phosphocreatine concentration estimates were corrected for the proportion of total brain matter within the VOI (i.e. divided by ([GM] + [WM])/([GM] + [WM] + [CSF])).

Reliability of total creatine (creatine + phosphocreatine, noted tCr) concentration estimate across all 4 measurements was assessed using intra-class correlations (two-way random effects model with absolute agreement) implemented in SPSS (SPSS inc., version 24.0). An excellent degree of test-retest reliability was observed for tCr (*ICC* = 0.86; *p* < 10^−3^), enabling us to use it for internal referencing. In all analyses, GABA and glutamate levels were therefore reported as ratios to tCr. Across the 2 baseline measurements, test-retest reliability was good for Glu:tCr (*ICC* = 0.75; *p* = 0.001) but lower for GABA:tCr (*ICC* = 0.28; *p* = 0.26). Thus, when evaluating the relationship between baseline metabolite concentration and computational parameters, we averaged the baseline concentration estimates across the 2 scanning sessions to obtain a more reliable estimate.

#### Diffusion-weighted images

Analysis of DW images was performed using tools from the FMRIB Software Library v5.0 (FSL; https://fsl.fmrib.ox.ac.uk) (Smith et al. 2004; Woolrich et al. 2009; Jenkinson et al. 2012). Preprocessing included the correction for susceptibility distortions, eddy current distortions and subject movement artefacts using TOPUP (https://fsl.fmrib.ox.ac.uk/fsl/fslwiki/topup) (Andersson et al. 2003) and EDDY (https://fsl.fmrib.ox.ac.uk/fsl/fslwiki/eddy) (Andersson and Sotiropoulos 2016). A diffusion tensor model was then fitted to the corrected DWI data using FDT (https://fsl.fmrib.ox.ac.uk/fsl/fslwiki/FDT) (Behrens et al. 2003; Behrens et al. 2007) in order to generate individual Fractional Anisotropy (FA) and Mean Diffusivity (MD) maps.

Voxelwise statistical analysis of the FA data was carried out using TBSS (Tract-Based Spatial Statistics, https://fsl.fmrib.ox.ac.uk/fsl/fslwiki/TBSS) (Smith et al. 2006b). In brief, all subjects’ FA and MD images were first brain-extracted using BET (https://fsl.fmrib.ox.ac.uk/fsl/fslwiki/BET) (Smith 2002) and then aligned into a common space using the nonlinear registration tool FNIRT (https://fsl.fmrib.ox.ac.uk/fsl/fslwiki/FNIRT) (Andersson et al. 2007). Next, the mean FA image was created and thinned to create a mean FA skeleton, which represents the centres of all tracts common to the group. Each subject’s aligned FA data was then perpendicularly projected onto this skeleton for statistical analysis. The same procedure was applied to MD data. Visual observation of the raw DWI data and skeletonised FA and MD maps revealed unreliable measurements in the temporal lobes due to large signal dropouts. Therefore, voxels from the temporal lobes were excluded from further statistical analysis.

The significance of the association between individual differences in weight of the fast and slow systems and white matter microstructure (FA and MD) was tested using Permutation Analysis of Linear Models (PALM) (https://fsl.fmrib.ox.ac.uk/fsl/fslwiki/PALM; version alpha-109) (Winkler et al. 2014; Winkler et al. 2016). The advantage of this technique is that it allows carrying out voxelwise joint inference on FA and MD using a full nonparametric permutation testing and Fisher combination to calculate Family-Wise Error (FWE)-corrected significance level. Results were considered significant for *p* < 0.05 after FWE 2D threshold-free cluster extent correction and correction across contrasts. The location of significant clusters was determined using the JHU DTI-based white matter atlases (https://fsl.fmrib.ox.ac.uk/fsl/fslwiki/Atlases) (Mori et al. 2005).

#### Resting state functional MRI

Analysis of resting state fMRI data was carried out using an independent component analysis (ICA) approach as implemented in MELODIC (https://fsl.fmrib.ox.ac.uk/fsl/fslwiki/MELODIC) (Beckmann et al. 2005) and tools from FSL (Smith et al. 2004). The first 5 volumes of each fMRI scan were discarded to allow for T1 equilibration effects. Standard pre-processing steps were applied, which included motion correction (MCFLIRT) (Jenkinson et al. 2002), brain extraction (BET) (Jenkinson et al. 2005), B0 unwarping (FUGUE), spatial smoothing using a Gaussian kernel of full-width at half-maximum (FWHM) of 5 mm, and high-pass temporal filtering equivalent to 100 s. fMRI volumes were registered to the individual’s structural scan using boundary-based registration (BBR) (Greve and Fischl 2009) and then to standard space images using FNIRT (Andersson et al. 2007). Individual fMRI data were denoised using FIX (https://fsl.fmrib.ox.ac.uk/fsl/fslwiki/FIX, version v1.065 beta) trained on a subset of 24 scans taken from our dataset (Griffanti et al. 2014).

Pre-processed functional data containing 280 volumes for each time point (4 time points per subject: pre/post, real/sham a-tDCS) and each subject (n = 22; Table 1) were temporally concatenated across subjects to create a single 4D data set that was decomposed into 25 components using ICA (Stagg et al. 2014; Bachtiar et al. 2015). All components were manually screened and non-physiological components were discarded. Informed decision was based on thresholded spatial maps (*Z* > 4, regional loci outside reasonable areas), as well as Fourier frequency decomposition of the components’ time courses (shift toward high frequencies) (Beckmann et al. 2005; Griffanti et al. 2014). In total, 13 components were kept (Figure 8). They included 3 “visual” networks (maps 1_25_, 2_25_ and 3_25_), the default-mode network (map 4_25_), an “auditory” network (map 5_25_), a “cerebellar” network (map 6_25_), an “insular” network (map 7_25_), an “executive control” network (map 8_25_), 2 “frontoparietal” networks (maps map 9_25_ and 10_25_) and 3 “sensorimotor” networks (maps 11_25_, 12_25_ and 13_25_). The 3 sensorimotor networks reflected the somatotopic organisation of the primary sensorimotor cortices and superior cerebellum, with a predominantly leg (medial), face (inferior) and hand (lateral) component (maps 11_25_, 12_25_ and 13_25_ respectively; Figure 8) (Van Den Heuvel et al. 2010).

**Figure 8:**
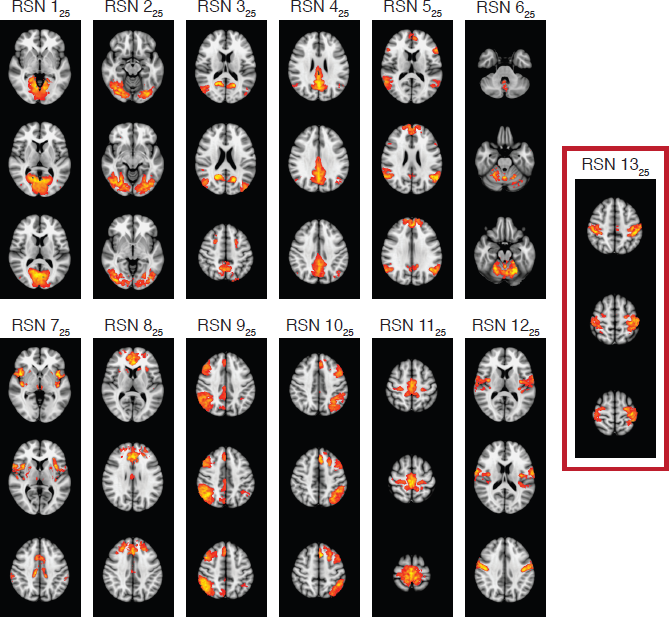
Thirteen resting-state networks extracted from the 25-component analysis. All ICA spatial maps were converted to z-statistic images via a normalised mixture-model fit and thresholded at *Z* > 4. The resting state network highlighted in a red square (RSN 13_25_) is the “hand” sensorimotor network and lies underneath the anode electrode.

Next, we estimated individual component spatial maps and corresponding time series for each subject and time point using dual regression (https://fsl.fmrib.ox.ac.uk/fsl/fslwiki/DualRegression) (Beckmann et al. 2009; Filippini et al. 2009). Network modelling was then carried out using the FSLnets toolbox (https://fsl.fmrib.ox.ac.uk/fsl/fslwiki/FSLNets) (Smith et al. 2015). Netmats were estimated using partial temporal correlations between components’ time series, with a small amount of L2 regularisation (ρ = 0.1 in the Ridge regression netmats option in FSLnets, i.e. netmats5). Pearson correlation coefficients were converted into Z-statistics with Fisher’s transformation before statistical analysis. The relationship between our computational parameters of interest (weights of the fast and slow systems) and the degree of functional connectivity between RSNs was investigated within a GLM framework. In order to minimise multiple comparisons, only edges involving the “hand” sensorimotor component (map 13_25_) were included in the analysis. Non parametric testing was used using PALM (Winkler et al. 2014; Winkler et al. 2016) with correction across contrasts.

## Appendix

### Statistical analysis of motor behaviour during and after prism adaptation

The online and offline experiments aimed to replicate the finding that left M1 a-tDCS would enhance AE consolidation when applied during but not before PA. A group of 24 participants underwent PA and tDCS in a repeated measures design, in 4 separate sessions, at least 1 week apart, in which they received anodal or sham tDCS either before or during PA. RM ANOVA considered the within-subjects of tDCS (anodal versus sham) and block separately for each experimental condition (online and offline).

- **Prism exposure (Blocks E1-7)** Accuracy on CLP improved progressively as adaptation developed in the online (Blocks E1-7: *F* (3.24; 74.44) = 132.24; *p* < 10^−3^; Figure 9A) and offline (Blocks E1-7: *F* (1.61; 2.21) = 89.51; *p* < 10^−3^; Figure 9B) experiments. No other effects were significant, indicating that neither online nor offline a-tDCS significantly influenced block mean pointing accuracy on CLP.
- **Adaptation phase (Blocks AE1-7)** Interleaved with exposure blocks, prism AE was measured as adaptation developed. Prism AEs stabilised progressively over time in both the online (Blocks AE1-7: *F* (3.95; 90.78) = 11.07; *p* < 10^−3^; Figure 9A) and offline (Blocks AE1-7: *F* (3.324; 76.456) = 15.408; *p* < 10^−3^; Figure 9B) experiments. However, no other effects were significant, indicating that neither online nor offline a-tDCS significantly influenced motor behaviour on OLP during the adaptation.
- **Retention phase (Blocks AE8-10)** Following PA and a 10 min rest period, AE was assessed. For the online experiment, a RM ANOVA with within-subjects factors of tDCS (anodal vs. sham) and blocks (AE8-10) confirmed the prediction of enhanced AE magnitude (main effect of tDCS: *F* (1; 23) = 4.38; *p* = 0.02, 1-tail, mean difference = −1.10 degrees, SEM = 0.53, Cohen’s *d* = 0.46; Figure 9A). As expected, this effect was not found in the offline experiment (*F* (1; 23) = 0.22; *p* = 0.64, 2-tail, mean difference = −0.294 degrees, SEM = 0.6221; Figure 9B).

**Figure 9:**
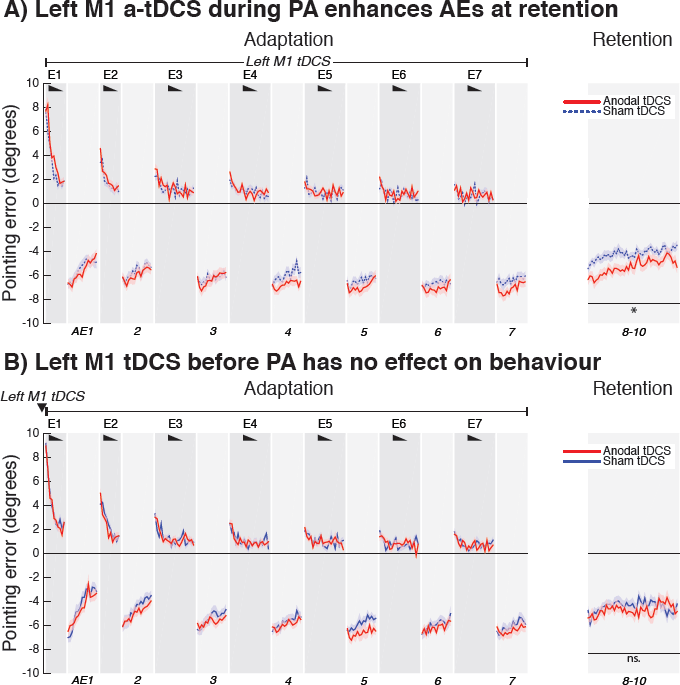
Behavioural effect of left M1 a-tDCS. The x-axis represents trial number and the y-axis pointing accuracy in reference to baseline accuracy (i.e. change from baseline). Positive values represent a rightward shift and negative values leftward shift. Pointing accuracy in healthy volunteers (n = 24) is plotted (group mean ± 1 SEM) when anodal (red) or sham (blue) was applied to the left M1 during PA (A) or before PA (B). Black wedges indicate blocks throughout which prisms were worn (CLP trials). During prism exposure (E1-7), participants saw the outcome of each trial, so they could gradually correct their errors. The AE was measured without visual feedback (AE1-7, shaded light grey). Similar to previous report (O’Shea et al. 2017), relative to sham, anodal tDCS increased prism AE at retention when applied during (*p* = 0.02; *Cohen’sd* = 0.46) but not before (*p* = 0.64). Asterisk indicates significant difference between the anodal and sham online condition (*p* < 0.05).

### Statistical analysis of the neurochemical effect of left M1 a-tDCS

RM ANOVAs considered the within-subject factors of time (pre vs. post tDCS) and tDCS (anodal vs. sham) separately for each metabolite to evaluate the neurochemical effect of the stimulation. At the group level, a-tDCS did not affect GABA (*F* (1; 15) = 1.10; *p* = 0.31), glutamate (*F* (1; 19) = 2.31; *p* = 0.15) or Glu:GABA (*F* (1; 15) = 0.25; *p* = 0.62) levels, as illustrated by no interaction ‘time × tDCS’ (Figure 10). In other words, at the group level, we found no evidence for the previously reported reduction in GABAergic inhibition with a-tDCS (Stagg et al. 2009; Bachtiar et al. 2015; Antonenko et al. 2017). Nevertheless, the neurochemical response to a-tDCS showed large inter-individual variability, which enabled us to address our question of interest: whether individuals with larger a-tDCS-induced increase in E:I ratio would also show larger increase in *R_w_* when a-tDCS is applied during PA (results reported above).

**Figure 10:**
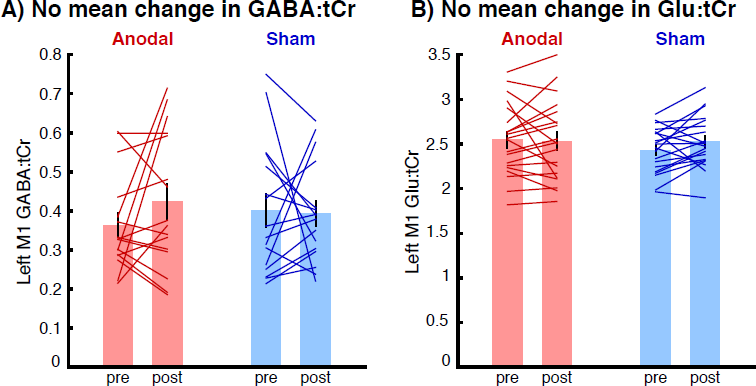
No group mean effect of stimulation on metabolite concentrations. Bar graphs represent the GABA:tCr ratio (A) and Glu:tCr ratio (B) within the MRS voxel before and after real (in red) or sham (in blue) left M1 a-tDCS inside the scanner. Individual participants are represented as lines overlaid on the bar graphs. Only individuals with complete dataset (i.e. satisfactory data quality for all 4 time points; see Table 1) are represented. RM ANOVAs revealed no significant interaction ‘time × tDCS’ for either metabolite (both *p* > 0.15). Note, however, the large inter-individual variability in neurochemical response to a-tDCS, exploited in Figure 4B.

### Statistical analysis of the a-tDCS effect on resting-state networks

Informed by the baseline relationship between *w_s_* and functional connectivity between the “hand” sensorimotor RSN and right frontoparietal RSN (edge 13_25_-9_25_), we asked whether the strength of this edge was influenced by M1 a-tDCS in the scanner. Repeated measure ANOVA revealed a main effect of scanning session (*F* (1; 21) = 5.70; *p* = 0.03) but no main effect of time (*F* (1; 21) = 0.10; *p* = 0.76) or interaction ‘time × stimulation’ (*F* (1; 21) = 0.55; *p* = 0.47). Additionally, inter-individual variations in the a-tDCS change in the edge 13_25_-9_25_ did not covary with inter-individual changes in *w_s_* (*r* = 0.08; *p* = 0.74, while controlling for the fast system). These null-results provide further support for the cognitive state dependency of the a-tDCS effect. The effect of a-tDCS on the functional coupling between the two networks is likely to require engaging individuals in the PA task, which was not the case inside the scanner.

A previous study reported increased functional connectivity within M1 with M1 a-tDCS (Bachtiar et al. 2015). In an attempt to replicate this result, we centred a 9 × 9 × 9 mm^3^ region of interest (ROI) on the peak coordinates of the “hand” sensorimotor RSN within the left M1 (x = −37, y = −24, z = 62; Z = 9.4) and extracted the mean network strength for each subject, stimulation condition, and time point. Unlike in Bachtiar et al. (2015)’s study, repeated measures ANOVA showed no significant main effect of stimulation (*F* (1; 21) = 0.39; *p* = 0.54) or time (*F* (1; 21) = 1.10; *p* = 0.31) and no significant stimulation x time interaction (*F* (1; 21) = 0.04; *p* = 0.84).

